# DNA-protein quasi-mapping for rapid differential gene expression analysis in non-model organisms

**DOI:** 10.1101/2022.12.15.520671

**Authors:** Kyle Christian L. Santiago, Anish M.S. Shrestha

## Abstract

**Background:** Conventional differential gene expression analysis pipelines for non-model organisms require computationally expensive transcriptome assembly. We recently proposed an alternative strategy of directly aligning RNA-seq reads to a protein database, and demonstrated drastic improvements in speed, memory usage, and accuracy in identifying differentially expressed genes.

**Result:** Here we report a further speed-up by replacing DNA-protein alignment by quasi-mapping, making our pipeline *>* 1000 × faster than assembly-based approach, and still more accurate. We also compare quasi-mapping to other mapping techniques, and show that it is faster but at the cost of sensitivity.

**Conclusion:** We provide a quick-and-dirty differential gene expression analysis pipeline for non-model organisms without a reference transcriptome, which directly quasi-maps RNA-seq reads to a reference protein database, avoiding computationally expensive transcriptome assembly.

## Background

Due to decreasing cost of sequencing, RNA-seq has become a mainstay of gene expression studies on a wide variety of organisms spanning the tree of life. A vast majority of these organisms do not have well-annotated reference genomes or transcriptomes, which are required by standard RNA-seq data analysis pipelines. A conventional work-around for such “non-model” organisms has been to assemble a transcriptome sequence from the RNA-seq reads, which is then used as reference. Functional inference of differentially expressed genes are done by aligning the assembled contigs to reference protein databases to find orthologs.

In some applications of RNA-seq in non-model organisms, transcriptome assembly, which requires massive computational resources, tends to be an overkill. This is especially the case when the main aim is to identify over or under-expressed genes, and not, say, to discover novel transcripts. In such cases there is no direct interest nor enough sequencing depth to reconstruct good quality transcript sequences. Apart from being resource hungry, transcriptome assembly is also known to introduce errors such as over-extension, mis-assembly, etc [1, 2].

In our prior work, we proposed the novel pipeline SAMAR [3], which directly aligns RNA-seq reads to a protein database – the one that would have been after-all used by the assembly-based approach for functional analysis – to measure expression and for subsequent differential expression analysis. Another recently proposed software tool Seq2Fun [4] also uses nucleotide-to-protein alignment for gene abundance quantification and functional profiling of RNA-seq data. An obvious outcome of this direct alignment approach is the drastic reduction of computational costs compared to the assembly-based approach. More interestingly, we showed that this approach has significantly higher precision and recall than assembly-based approach in identifying differentially expressed genes.

Here we sought to further speed up SAMAR, by exploiting the fact that gene expression levels can be estimated based on just the knowledge of which reference sequence(s) each read maps to, without the need to compute alignments. For the problem of mapping RNA-seq reads to a reference transcriptome, this idea of replacing traditional alignment by faster mapping techniques is known to provide significant speed-ups [5, 6, 7]. We adapted to our case of mapping RNA-seq reads to a reference protein database, one such mapping technique, called quasi-mapping, the main idea of which is to determine the mapping by rapid look-ups of sub-strings of a query sequence. We incorporated our quasi-mapping technique into a pipeline for differential analysis. We show that our pipeline is *>* 1000× faster while remaining to be more accurate than assembly-based approach. Compared against other alignment/mapping strategies, we show that our method is fastest at the cost of sensitivity.

An implementation of our pipeline SAMAR-lite is available at https://bitbucket.org/project_samar/samar_lite.

## Methods

### Notations

For a string *S*, we use *S*[*i* : *j*] to denote the substring of *S* starting at position *i* and ending at position *j*. We use |*S*| to denote the length of *S*.

### Reference index construction

In the first stage of our method, we construct an index of the reference set of proteins to allow quick sub-string searches. We describe the data-structures below with the aid of an example shown in Figure 1. We concatenate the reference sequences into a string *C*, separating the individual sequences by the special character $. We construct the suffix array *SA* of *C*, which is an array containing the indices of the sorted suffixes of *C*, and allows for fast sub-string searches using binary search. We augment *SA* with a hash table *HT* which maps a k-mer *S* present in *C* to *HT* (*S*) which is an interval in *SA* containing suffixes that have *S* as prefix. The role of *HT* is to allow faster searches in *SA* by reducing the search area of binary search. This idea has been explored in greater detail in [8]. Additionally, we construct a bit vector *B* of the same length as *C* containing 1 at each position corresponding to the special character $ and 0 elsewhere. A rank query *rank*_*B*_(*i*) takes a position *i* of *SA* as input and returns the number of 1s up to position *i* in *B*, essentially identifying the protein sequence in *C* corresponding to position *i* in the suffix array. A rank query can be accomplished in constant time by representing *B* using a succinct data-structure. A rank query can be generalized to *rank*_*B*_([*i, j*]), where it takes an interval [*i, j*] of *SA* and returns the corresponding set of proteins.

**Figure 1.**
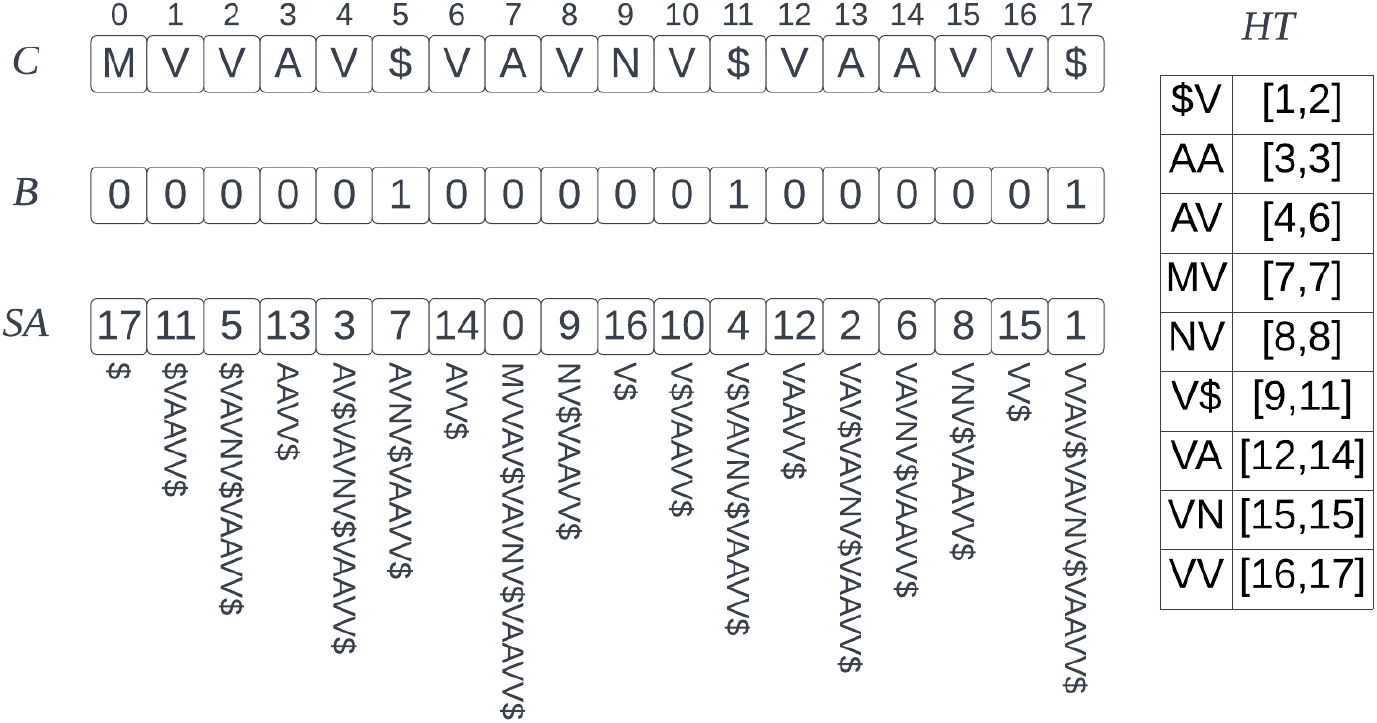
Example of the reference index data-structures for the set of sequences MVVAV, VAVNV, VAAVV. *C* is the concatenated string, *B* is the bit-vector, *SA* suffix array, and *HT* is a hash table mapping a 2-mer to a suffix array interval containing suffixes whose prefix is the 2-mer.

### Quasi-mapping

In the second stage, we map each RNA-seq read to protein sequence(s) likely to be the translation product of the transcript from which the read originates. Our method closely follows the quasi-mapping technique of [6]. The core idea is to determine the mapping based on the set of k-mers and extended k-mer matches that are found rapidly using the data-structures described above.

In more detail, for each read, we first translate it into six amino acid sequences, one for each possible reading frame. Let *T* be one such translation. We iterate through the positions of *T*, performing at a position *pos*, the actions described below and also in the psuedo-code in Algorithm 1. If the k-mer starting at *pos* is a key in *HT*, we obtain its corresponding suffix array interval *sai*. We next attempt to extend the match by adding the amino acid at *pos* + *k* to the current k-mer and performing a binary search within *sai*. We repeat this extension process until no matches can be found. At the end of the extension process, let *k*^*′*^ be the length of the extended k-mer and *sai*^*′*^ be the maximal suffix array interval containing suffixes whose prefix is *T* [*pos* : *pos* + *k*^*′*^]. Let ℙ be the set of protein sequences whose sub-strings are in the interval *sai*^*′*^. For each *P* ∈ ℙ, we record that there was a *k*^*′*^-length substring match. The next iteration begins at *pos* + *k*^*′*^, or at *pos* + 1 if the k-mer at *pos* was not in *HT* to begin with.

This mapping procedure is run six times, once for each possible reading frame. As the final mapping output, we return all protein sequences whose coverage, which is the total length of sub-string matches found, is larger than a pre-specified threshold, for any of the reading frames.

### Counting

The mappings generated from the previous step are used to measure expression levels to be used for differential expression analysis. As in SAMAR, we follow the counting strategy of rescuing multi-mapping reads [9]. The counting is done in two passes. First, for each sequence *P* in the reference, we count the number of reads mapping uniquely to it, and normalize the count by |*P* |. Let this normalized count be *n*_*P*_. Second, for each multi-mapping read, we update the count of each sequence *P* it multi-maps to, in proportion to *n*_*P*_. That is, we add to *n*_*P*_ the value *n*_*P*_ */* ∑ *n*_*Q*_, where the summation is over all *Q*s to which the read maps. If the denominator is zero, we distribute the count evenly among the proteins the read multi-maps to.

#### Algorithm 1: Quasi-mapping of one translation of a read. This process is repeated for all possible 6 frames

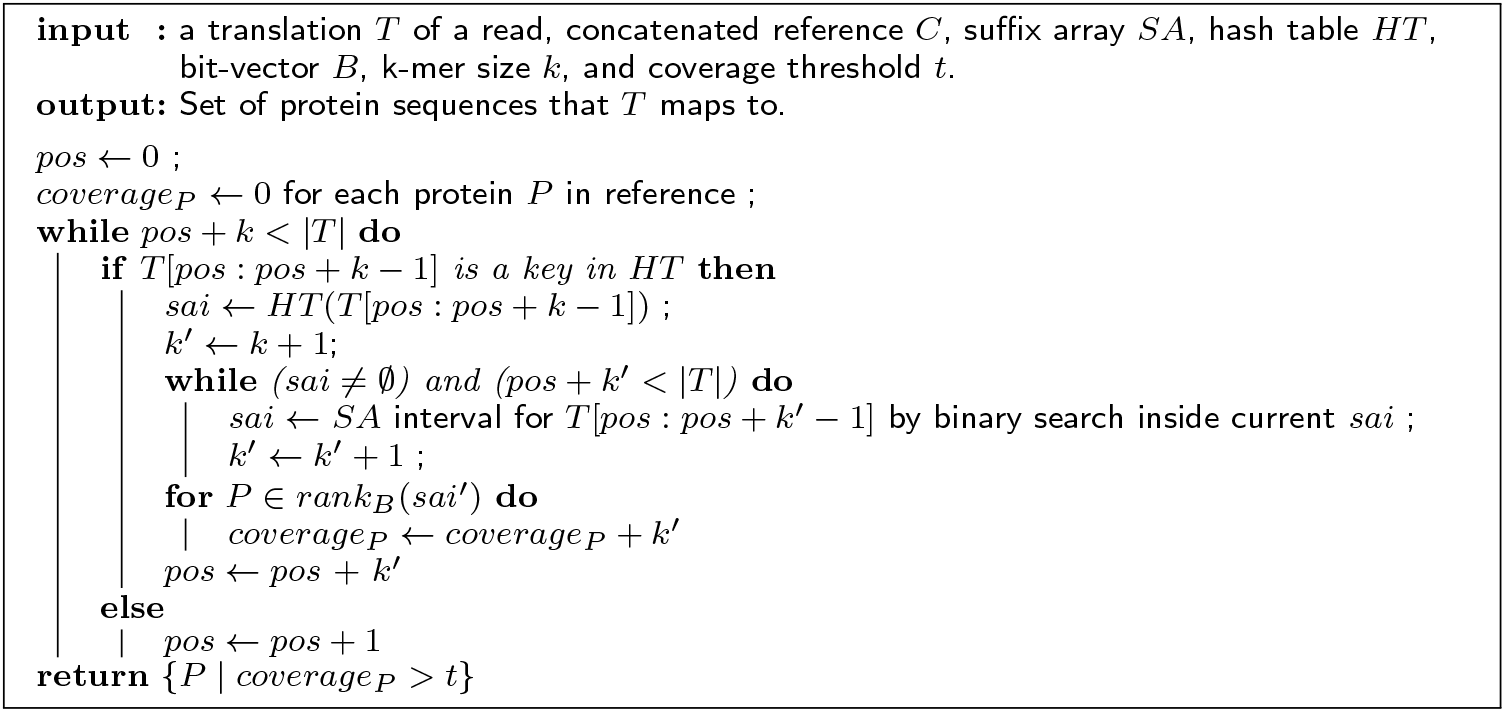

### Implementation details

We implemented the quasi-mapping method in the Rust programming language [10] using the RustBio library [11] for the suffix array and succinct representation of the bit vector for efficient rank queries. We incorporated the mapping step, the counting step, and DESeq2 [12] for differential analysis into a Snakemake [13] pipeline. The software is available at https://bitbucket.org/project_samar/samar_lite.

## Results

### Read simulation

For benchmarking, we used an RNA-seq read dataset that simulates a typical small-sample-size and low-coverage whole-genome gene expression experiment from our prior work [3]. The dataset was generated using Polyester [14] from the protein-coding transcriptome (BDGP6.28) of *D. melanogaster* obtained from Ensembl Genes 101 and containing 28,692 transcripts of 13,320 genes. We simulated two groups of read sets, with three replicates in each group. For the first group, we set the mean expression levels to be proportional to the FPKM values from an arbitrary poly-A+ enriched real RNA-seq data (ArrayExpress E-MTAB-6584). In the second group, around 30% of the transcripts were simulated to be differentially expressed with different levels of up-regulation and down-regulation. The transcripts were chosen by randomly selecting genes and setting only the highest expressing isoform to be differentially expressed. Each read set had roughly 20 million pairs of 100 bp reads with mean fragment length of 250 bp.

### Mapping performance and run time

#### Reference proteomes used

We first evaluated the performance of our method in the task of mapping the reads to four reference proteomes at varying levels of evolutionary divergence from the read source. The *D. melanogaster* proteome (Uniprot ID UP000000803) provides a baseline for performance comparison. The proteomes of *D. ananassae* (UP000007801) and *D. grimshawi* (UP000001070) were used to study the effect of mapping to the proteome of close relatives. Additionally, the proteome of *Anopheles gambiae* (UP000007062) was used to study the effect of mapping to a distant relative. Phylogenetic studies place *D. melanogaster* closer to *D. ananassae*, with a separation of 10-20 MY, than *D*.*grimshawi*, with separation of around 40 MY (see e.g. [15, 16]). According to one estimate, the lineages of *D. melanogaster* and *A. gambiae* separated roughly 250 MYA, and the average sequence identity at amino acid level in orthologous exons is only around 61.6% [17].

#### Tools compared

We compared our mapping performance against several existing tools for DNA-protein alignment/mapping of large data: LAST [18] (version 1060), DIAMOND [19] (version 2.0.12), and Kaiju [20] (version 1.9.0). We selected LAST as it was the alignment tool used in SAMAR [3]. We selected DIAMOND because it is a widely used DNA-protein alignment tool for big sequence data, although mainly focusing on metagenomic data. Kaiju, which employs maximum exact matches (MEM) to compute mappings, was originally proposed for metagenomic data, but it has recently been used by Seq2Fun [4] for RNA-seq dataset profiling. The commands used to run each tool is provided in the Supplementary Material.

#### Evaluation metric

We evaluated mapping correctness as follows. Consider a read *r* from a transcript of *D. melanogaster* gene *g*. Let *M*_*g*_ be the set of protein products of *g*. When using *D. melanogaster* proteome as reference, *r* was defined to be correctly mapped if at least one of the proteins it was mapped to, is in *M*_*g*_. When using the other references, we determined correctness using pre-computed orthology maps between *D. melanogaster*–*D. ananassae* and *D. melanogaster*–*D. grimshawi* and *D. melanogaster* – *A. gambiae* pairs, which were obtained from InParanoid [21]. We defined *r* as being correctly mapped if at least one of the proteins it was mapped to is in an ortholog group containing a protein from *M*_*g*_. A read with no mappings reported is not counted as correct or incorrect.

Mapping performance and running times are plotted in Figure 2 and Figure 3, respectively.

**Figure 2.**
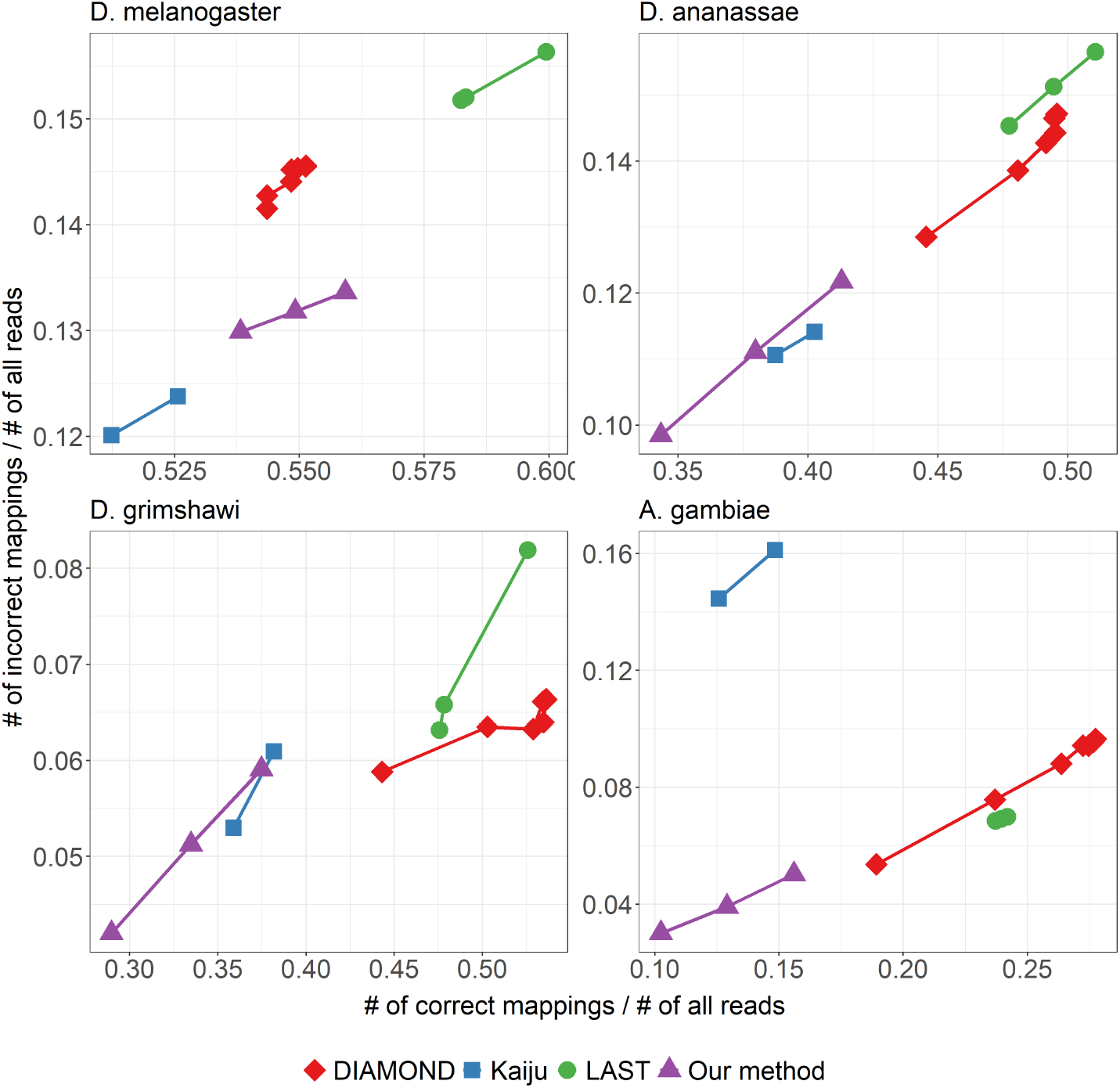
Mapping performance of the different aligners/mappers when using different reference proteomes. For DIAMOND, different points correspond to the different available presets: normal, fast, mid-sensitive, sensitive, more-sensitive, very-sensitive, and ultra-sensitive. For Kaiju, the points correspond to setting the mode to greedy and MEM. For LAST, the maximum mismap probability was set to 0.95,0.9, and 0.8. For our method, we used the k-mer size of 7 and coverage threshold was varied among 40, 50, and 60.

**Figure 3.**
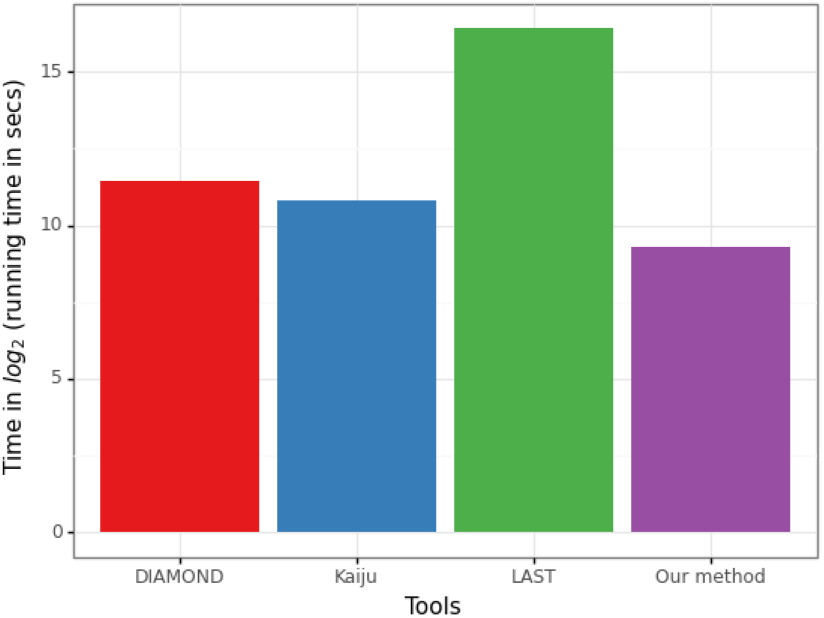
Comparison of mapping run times for a typical sample containing roughly 21 million pairs reads of length 100 bp each. Each tool was run on a single thread and with the following settings: normal for DIAMOND, greedy for Kaiju, maximum mismap probability of 0.95 for LAST, and k-mer 7 with coverage threshold 40 for our method

### Differential expression analysis performance

Next, we fed the alignments/mappings produced by each tool to downstream differential expression analysis and compared their performance in identifying differentially expressed genes.

#### Pipelines compared

Of the possible parameters for each tool, we used the normal preset for DIAMOND, greedy for Kaiju, maximum mismap probability of 0.95 for LAST, and *k*-mer 7 with coverage threshold 40 for our method. The mappings were counted using the method described in the Methods section. For LAST, we used a slightly different variation offered in SAMAR, in which the counts are normalized by the length of the regions that have reads mapping to them rather than the entire length of the protein. The counts are then passed to DESeq2 [12] for differential expression analysis.

Additionally, to mimic the case of a non-model organism, we pretended that *D. melanogaster* reference transcriptome doesn’t exist; and we ran a typical assembly-based pipeline consisting of: Trinity [22] for de-novo transcriptome assembly, followed by Bowtie2 [23] for mapping the reads to the assembly, RSEM [24] for counting, tximport [25] for gene-level aggregation using the gene-to-transcript mapping provided by Trinity, and finally DESeq2 [12] for differential analysis. Dammit was used for annotating the assembled contigs against the *D. ananassae* and *D. grimshawi* proteomes. Since the annotation process produces many short local alignments, we keep only those alignments covering at least 50% of the length of the contig as evidence of homology. Commands used for assembly are described in the Supplementary Material.

#### Evaluation metric

We evaluated each pipeline based on their precision and recall in predicting differentially expressed genes. Let *M*_*g*_ be the set of protein products of a *D. melanogaster* gene *g*. When using the *D. melanogaster* reference, an actual up-regulated (down-regulated) gene *g* is defined as correctly predicted if at least one protein in *M*_*g*_ (defined in previous section) was predicted to be up-regulated (down-regulated). When using the proteome of a relative as reference, *g* is defined as correctly predicted if at least one protein in an ortholog group containing *M*_*g*_ was predicted to be up-regulated (down-regulated). We define precision as the proportion of predicted differentially expressed genes that are actually differentially expressed, and recall as the proportion of actual differentially expressed genes that were correctly predicted to be differentially expressed.

The results are shown in Figure 4.

**Figure 4.**
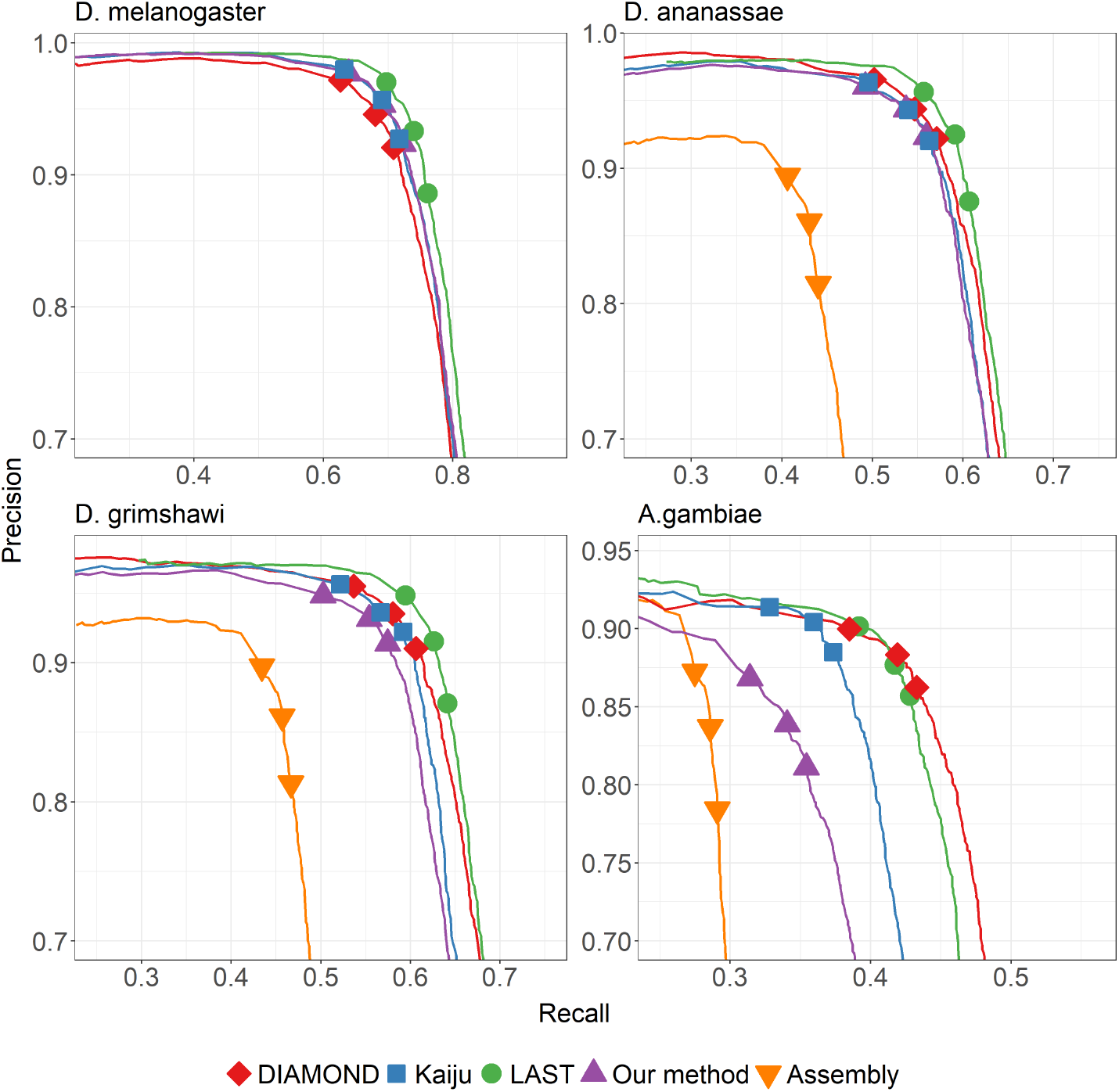
Precision-recall curves for differential gene expression analysis using different tools for the mapping step and when using different reference proteomes. The three shape markers in each curve correspond to setting the false discovery rates in DESeq2 to the values of 0.01, 0.05, and 0.1.

## Discussion

### Our method is *>* 1000× faster than assembly-based approach while being more accurate

We set out to build a quick-and-dirty differential gene expression analysis pipeline by avoiding transcriptome assembly and replacing a slower alignment step with quasimapping. As can be seen from the *D. grimshawi* and *D. ananassae* plots in Figure 4, even when replacing sensitive full alignment step with the quasi-mapping step, our pipeline for identifying differentially expressed genes outperforms assembly-based approach. This gain comes with dramatic speed-up. Transcriptome assembly of a dataset containing a total of roughly 120 million pairs of reads, with 4 cores and 8 threads engaged took more than 7 days to complete. Our mapping takes less than 10 minutes for handling the same data, making it more than 100 times faster.

### When compared to other aligners/mappers, our method provides a trade-off between speed and sensitivity

As seen in the run-time plot of Figure 3, our method runs the fastest among all the tools – being *>* 2.5× faster than the next fastest Kaiju, *>* 4× faster than DIAMOND, and *>* 100× faster than LAST. This might not be surprising as DIAMOND and LAST compute alignments using seed-and-extend approach, and LAST additionally computes appropriate alignment scoring scheme as well as alignment column probabilities. On the other hand, our method and Kaiju rely on finding exact matches. However, our speed-up comes at the cost of lesser mapping accuracy. As can be seen in Figure 4, for the case of using the proteome of close relatives *D. ananassae* or *D. grimshawi* as reference, our method is overall less sensitive and precise than LAST and DIAMOND, while performing roughly similar as Kaiju.

When using the proteome of the distant relative *A. gambiae*, the performance of our method worsens to the greatest degree compared to other aligners/mappers. The results of Figure 4 essentially capture the differences in mapping performance shown in Figure 2. There is a stark performance gap compared to LAST and DIAMOND, with our method correctly mapping 10-15% fewer reads than LAST or DIAMOND. Compared to Kaiju, the difference in mapping performance is not too apparent for close relatives, but Kaiju performs better when the reference becomes more distant.

We note that while we chose various reference proteomes to demonstrate the effect of varying levels of evolutionary divergence, a part of the differences we see in Figures 2 and 4 might be attributed to the differences in the assembly and annotation pipeline employed to generate those reference sequences.

### Reduced alphabet

We implemented quasi-mapping on the reduced amino-acid alphabet proposed by DIAMOND: *{*K,R,E,D,Q,N}, {C}, {G}, {H}, {I,L,V}, {M}, {F}, {Y}, {W}, {P}, {S,T,A}, where characters in the same set are treated to be equivalent. The results are shown in Figure 5. We observed that for the reduced alphabet, coverage threshold value of 40 – corresponding to rightmost points in each curve – result in a substantial increase in incorrect mappings compared to the non-reduced alphabet. At the coverage threshold of 50, *k* = 11 for reduced alphabet is close to the performance of *k* = 7 for non-reduced alphabet, while being almost 2.5× faster.

**Figure 5.**
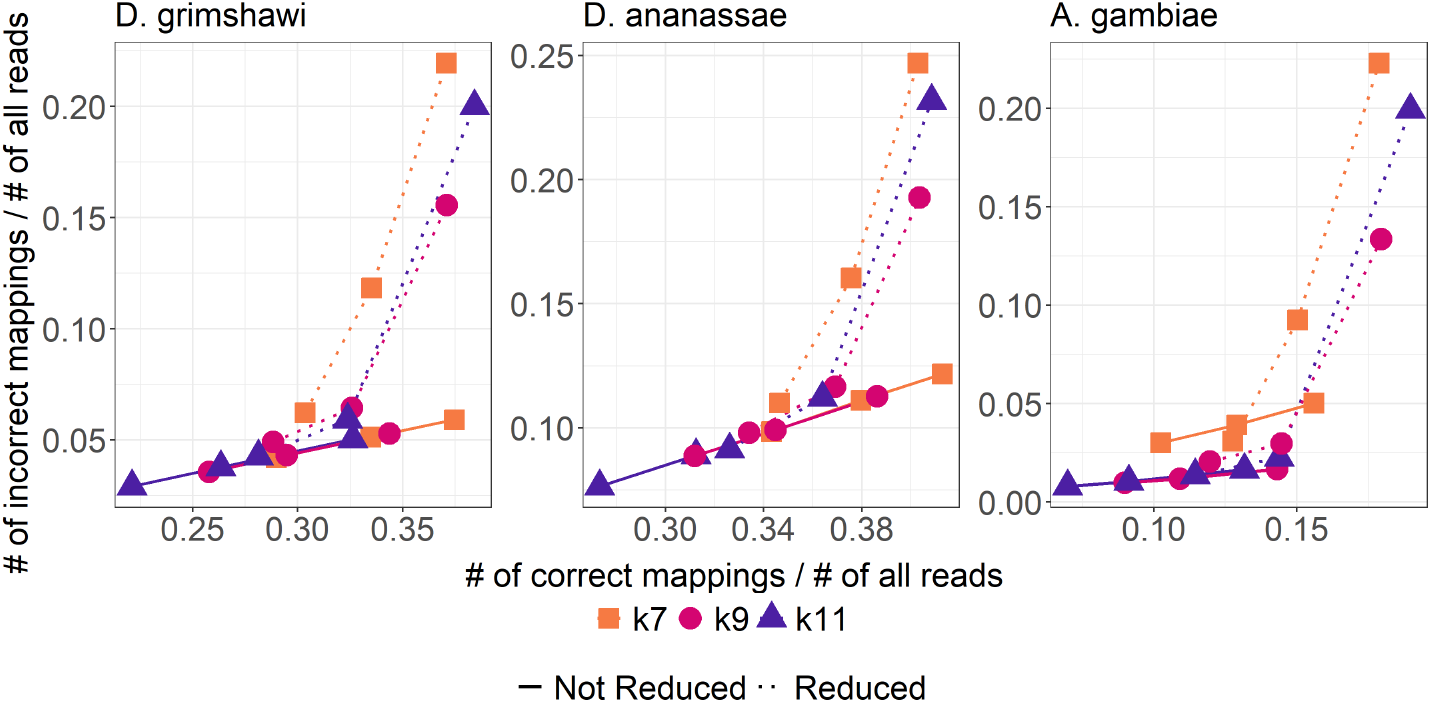
Comparing the performance of our method on the full amino acid alphabet versus a reduced one of size 11. For each curve, the three points from left to right correspond to coverage thresholds of 60, 50, and 40, respectively.

### Effect of reference proteome

The availability of an accurate reference protein database is key to the performance of all methods evaluated in this paper, but more so for our method which relies on exact k-mer matches. This is seen, perhaps unsurprisingly, in our evaluations, where our method seems to be most sensitive to evolutionary divergence. Another factor that could affect performance is the presence of large number of highly similar sequences in the reference. This could be as a result of high level of recent gene/genome duplications, high amount of repetitive sequences in transcripts due to transposable elements, or because the reference spans multiple species and contains orthologs (e.g. UniProtKB). For all methods, it would be interesting to investigate the extent of performance degradation due to these factors at the level of mapping and in downstream functional analysis. As a practical remedy for the case of multi-species reference, it might be better to first reduce redundancy by running sequence clustering tools (e.g. CD-HIT [26] or MMSeq [27]) or find orthogroups (e.g. using OrthoFinder [28]).

### Future directions

It would be interesting to reduce the gap in sensitivity compared to traditional seed-and-extend methods without raising computational cost. Some promising direction include using spaced k-mers [29], syncmers [30], and minimally overlapping words [31]. Using spaced k-mers, for example, has been shown to be more sensitive than contiguous ones in other alignment-free sequence comparison applications [32], and are in fact also implemented in DIAMOND and LAST.

We also note that we chose the augmented suffix array data structure with memory usage not in mind, especially since our method uses negligible memory compared to de-novo transcriptome assembly. It might be interesting to explore other compact text indexes for cases where memory requirement is a concern.

## Conclusion

We have implemented a quick-and-dirty differential gene expression analysis pipeline for non-model organisms without a reference transcriptome. It uses quasi-mapping to rapidly map RNA-seq reads to a reference protein database, followed by a simple counting step the result of which can be fed to standard differential analysis tools like DESeq2. It is computationally super light-weight compared to the conventional approach of first building a transcriptome assembly. The quasi-mapping step itself is fast compared to other alignment/mapping techniques, but there is room for improvement in its sensitivity.

## Declarations

### Ethics approval and consent to participate

Not applicable.

### Consent for publication

Not applicable.

### Availability of data and materials

Our software is available at https://bitbucket.org/project_samar/samar_lite.

Datasets for benchmarking were obtained from public repositories: transcripts of protein-coding genes were obtained from the fruit fly assembly BDGP6.28 in Ensembl Genes 101; reference proteomes of *D. melanogaster* (UP000000803), *D. grimshawi* (UP000001070), *D. ananassae* (UP000007801), and *A. gambiae* (UP000007062) were obtained from UniProt.

### Author’s contributions

AS planned the study. KS wrote the software. AS and KS performed the benchmarking study. All authors wrote, read, and approved the manuscript.

## Acknowledgements

Kyle Santiago is thankful to the Department of Science and Technology and the Engineering Research and Development for Technology (ERDT) scholarship program for funding his Master of Science in Computer Science.

## Additional Files

Additional file 1 — Supplementrary Material

## References

1. Vijay, N., Poelstra, J.W., Künstner, A., Wolf, J.B.W.: Challenges and strategies in transcriptome assembly and differential gene expression quantification. a comprehensive in-silico assessment of RNA-seq experiments. Molecular Ecology 22(3), 620–634 (2012)

2. Hsieh, P.-H., Oyang, Y.-J., Chen, C.-Y.: Effect of de novo transcriptome assembly on transcript quantification. Scientific Reports 9(1) (2019)

3. Shrestha, A.M.S., Guiao, J.E.B., Santiago, K.C.L.: Assembly-free rapid differential gene expression analysis in non-model organisms using DNA-protein alignment. BMC Genomics 23(1) (2022)

4. Liu, P., Ewald, J., Galvez, J.H., Head, J., Crump, D., Bourque, G., Basu, N., Xia, J.: Ultrafast functional profiling of RNA-seq data for nonmodel organisms. Genome Research 31(4), 713–720 (2021)

5. Patro, R., Duggal, G., Love, M.I., Irizarry, R.A., Kingsford, C.: Salmon provides fast and bias-aware quantification of transcript expression. Nature Methods 14(4), 417–419 (2017)

6. Srivastava, A., Sarkar, H., Gupta, N., Patro, R.: RapMap: a rapid, sensitive and accurate tool for mapping RNA-seq reads to transcriptomes. Bioinformatics 32(12), 192–200 (2016)

7. Bray, N.L., Pimentel, H., Melsted, P., Pachter, L.: Near-optimal probabilistic RNA-seq quantification. Nature Biotechnology 34(5), 525–527 (2016)

8. Grabowski, S., Raniszewski, M.: Compact and hash based variants of the suffix array. Bulletin of the Polish Academy of Sciences Technical Sciences 65(4), 407–418 (2017)

9. Mortazavi, A., Williams, B.A., McCue, K., Schaeffer, L., Wold, B.: Mapping and quantifying mammalian transcriptomes by RNA-seq. Nature Methods 5(7), 621–628 (2008)

10. Matsakis, N.D., Klock, F.S.: The rust language. ACM SIGAda Ada Letters 34(3), 103–104 (2014)

11. Köster, J.: Rust-bio: a fast and safe bioinformatics library. Bioinformatics 32(3), 444–446 (2015)

12. Love, M.I., Huber, W., Anders, S.: Moderated estimation of fold change and dispersion for RNA-seq data with DESeq2. Genome Biology 15(12) (2014)

13. Mölder, F., Jablonski, K.P., Letcher, B., Hall, M.B., Tomkins-Tinch, C.H., Sochat, V., Lee, S., Twardziok, S.O., Kanitz, A., Wilm, A., Holtgrewe, M., Rahmann, S., Nahnsen, S., Köster, J.: Sustainable data analysis with snakemake. F1000Res 10(33) (2021)

14. AC, F., AE, J., B, L., JT., L.: Polyester: simulating rna-seq datasets with differential transcript expression. Bioinforma 31, 2778–2784 (2015)

15. Bhutkar, A., Russo, S.M., Smith, T.F., Gelbart, W.M.: Genome-scale analysis of positionally relocated genes. Genome Research 17(12), 1880–1887 (2007)

16. Haubold, B., Pfaffelhuber, P.: Alignment-free population genomics: An efficient estimator of sequence diversity. G3 Genes|Genomes|Genetics 2(8), 883–889 (2012)

17. Bolshakov, V.N., Topalis, P., Blass, C., Kokoza, E., della Torre, A., Kafatos, F.C., Louis, C.: A comparative genomic analysis of two distant diptera, the fruit fly, drosophila melanogaster, and the malaria mosquito, anopheles gambiae. Genome research 12, 57–66 (2002)

18. Yao, Y., Frith, M.C.: Improved DNA-versus-protein homology search for protein fossils. In: Algorithms for Computational Biology, pp. 146–158. Springer, ??? (2021)

19. Buchfink, B., Xie, C., Huson, D.H.: Fast and sensitive protein alignment using DIAMOND. Nature Methods 12(1), 59–60 (2014)

20. Menzel, P., Ng, K.L., Krogh, A.: Fast and sensitive taxonomic classification for metagenomics with kaiju. Nature Communications 7(1) (2016)

21. Sonnhammer, E.L.L., Östlund, G.: Inparanoid 8: orthology analysis between 273 proteomes, mostly eukaryotic. Nucleic Acids Research 43(D1) (2014)

22. Haas, B.J., Papanicolaou, A., Yassour, M., Grabherr, M., Blood, P.D., Bowden, J., Couger, M.B., Eccles, D., Li, B., Lieber, M., MacManes, M.D., Ott, M., Orvis, J., Pochet, N., Strozzi, F., Weeks, N., Westerman, R., William, T., Dewey, C.N., Henschel, R., LeDuc, R.D., Friedman, N., Regev, A.: De novo transcript sequence reconstruction from rna-seq using the trinity platform for reference generation and analysis. Nature Protocols 8, 1494–1512 (2013)

23. Langmead, B., Trapnell, C., Pop, M., Salzberg, S.L.: Ultrafast and memory-efficient alignment of short dna sequences to the human genome. Genome Biology 10(R25) (2009)

24. Li, B., Dewey, C.N.: Rsem: accurate transcript quantification from rna-seq data with or without a reference genome. BMC Bioinformatics 12(323) (2011)

25. Soneson, C., Love, M.I., Robinson, M.D.: Differential analyses for rna-seq: transcript-level estimates improve gene-level inferences. F1000Research 4(1521) (2015)

26. Li, W., Godzik, A.: Cd-hit: a fast program for clustering and comparing large sets of protein or nucleotide sequences. Bioinformatics 22(13), 1658–1659 (2006)

27. Steinegger, M., Söding, J.: Clustering huge protein sequence sets in linear time. Nature Communications 9(1) (2018)

28. Emms, D.M., Kelly, S.: OrthoFinder: phylogenetic orthology inference for comparative genomics. Genome Biology 20(1) (2019)

29. Ma, B., Tromp, J., Li, M.: PatternHunter: faster and more sensitive homology search. Bioinformatics 18(3), 440–445 (2002)

30. Edgar, R.: Syncmers are more sensitive than minimizers for selecting conserved k-mers in biological sequences. PeerJ. 9 (2021)

31. Frith, M.C., Noé, L., Kucherov, G.: Minimally-overlapping words for sequence similarity search. Bioinformatics 36(22–23), 5344–5350 (2020)

32. Boden, M., Schöneich, M., Horwege, S., Lindner, S., Leimeister, C.-A., Morgenstern, B.: Alignment-free sequence comparison with spaced k-mers. German Conference on Bioinformatics 2013 (2013)

