## Supplementary Material for "DNA-protein quasi-mapping for rapid differential gene expression analysis in non-model organisms"

### A Commands used for benchmarking aligners/mappers

All aligners/mappers were run on a system with an Intel i7-4710 CPU with 4 cores and 8 threads, and 32GBytes of RAM.

#### A.1 Our method (SAMAR-lite)

The reference index was constructed using:

```
ref-align reference.fa 7
```

where 7 is the k-mer size. This was followed by mapping with a specified coverage threshold, e.g. 40 below :

```
alignr reference_7.json query.fastq 40 output
```

This is followed by counting:

```
python seq_count.py reference.fa input output
```

This ends with DESeq2, which performs differential expression analysis on the counts:

```
txi <- tximport(files_counts, type = "salmon", txIn = TRUE, txOut = TRUE)
ourMethod_dds <- DESeqDataSetFromTximport(txi, colData = colData,
                                          design = ~ condition)
ourMethod_dds <- DESeq(ourMethod_dds)
```

#### A.2 LAST 1060

The reference index was constructed using:

```
lastdb -p index reference.fa
```

This was followed by aligment scoring scheme training:

```
last-train --revsym --matsym --gapsym --sample-number=600000 \
    -S0 index sample.fa > train_output
```

where `sample.fa` contains 6 amino-acid sequences (one each for a translation frame) per read of a sample of the input reads in fastq format.
This was followed by alignment:

```
lastal -p train_out -F20 index translated_reads.fa | \
      last-pair-probs -f mean -s std -m 0.95 -d 0.1 | \
      maf-convert tab" > output
```

This was followed by counting taken from the SAMAR [?] pipeline and DESeq2.

#### A.3 DIAMOND 2.0.12

The reference index was constructed using:

```
diamond makedb --in reference.fa -d reference
```

This was followed by the alignment:

```
diamond blastx -q query.fa -d reference -o out.tsv
```

This was followed by counting, similar to the counting of our method, and DESeq2.

#### A.4 Kaiju 1.9.0

The reference index was constructed using:

```
kaiju-mkbwt -n 8 -a ACDEFGHIKLMNPQRSTVWY -o reference reference.fa

kaiju-mkfmi reference
```

Where `n` is the number of threads, `a` is the alphabet, and `o` is the output Burrows-Wheeler transform to be converted to an FM-index in the next command.

This was followed by mapping:

```
kaijux -f reference -i query.fastq -o output.kaiju
```

This was followed by counting, similar to the counting of our method, and DESeq2.

### B Commands used for benchmarking an assembly-based pipeline

#### Assembly-based approach

De-novo transcriptome assembly was computed from all the reads in the dataset using Trinity (version 2.8.5):

```
Trinity --seqType fa --max_memory 32G --left reads1_1.fa,..,reads6_1.fa \
        --right reads1_2,..,reads1_6.fa --CPU 8
```

The reads were aligned to the assembled transcripts using Bowtie2 (version 2.4.1):

```
bowtie2-build Trinity.fasta index
bowtie2 -p 20 -x index -1 reads1_1.fa -2 reads1_2.fa --no-mixed \
     --no-discordant --gbar 100 --end-to-end -k 200 | \
     samtools view -bS - > alns.bam
```

Counting was done using RSEM:

```
rsem-prepare-reference Trinity.fasta rsem_index
rsem-calculate-expression -q --no-bam-output --alignments \
                        --paired-end alns.bam rsem_index counts
```

Transcript-level counts were aggregated at the gene level using tximport, based on the gene-transcript map constructed by Trinity, and finally differential expression analysis was performed using DEseq2.

```
contig2gene <- read.delim("Trinity.fasta.gene_trans_map", header=FALSE)[,c(2,1)]
txi <- tximport(files_counts, type = "rsem", txIn = TRUE, txOut = FALSE,
                tx2gene = contig2gene)
```

```
assembly_dds <- DESeqDataSetFromTximport(txi, colData = colData,
                                         design = ~ condition)
assembly_dds <- DESeq(assembly_dds)
```

Annotation was done against the reference proteome using Dammit.

```
dammit annotate Trinity.fasta --quick --user-databases dro_me_ref.fa -e 1e-10
```

Of the alignments that were reported in `Trinity.fasta.x.dro\_me\_ref.fa.crbl.csv}`, we kept only those that cover at least 50% of the contig length.
